# The Clash of Academic Hierarchy and Inclusive Leadership: Evolution of Leadership in a Nationwide Diversity, Equity, and Inclusion Initiative

**DOI:** 10.1101/2023.03.12.532284

**Authors:** Diane Codding, Bennett B. Goldberg

**Author notes:** Correspondence concerning this article should be addressed to Diane Codding, Northwestern University, 2145 Sheridan Road, Evanston, IL 60208. **Author Note** We acknowledge the support of members of the Alliance in guiding, reviewing, and encouraging this scholarly analysis and presentation of leadership transition. We acknowledge that for some, bravery in the face of revisiting trauma was required, and we very much appreciate being provided this opportunity. The authors report there are no competing interests to declare.

## Abstract

Leadership has been traditionally envisioned and enacted as a top-down structure that reinforces positionality and power dynamics that are antithetical to the fundamental values and desired outcomes of equity-focused change work. This paper critically examines the evolution of leadership structures, agenda setting, and decision-making within a large, multi-institution, federally funded, and equity-focused alliance. Findings from this case study suggest that developing and sustaining inclusive leadership structures requires significant resources and an enduring commitment to engaging in a critically reflexive process to redress inequities within equity-focused initiatives.

---

Leadership is arguably the central defining aspect of any organization or collective purposely brought together for specific outcomes. Kruse (2013) defines leadership as “a process of social influence, which maximizes the efforts of others, towards the achievement of a goal.” This process of social influence is embodied by the leadership structures that organizations and their leaders choose to implement. Thus, leadership structures can be understood as a functional expression of group values, norms, and culture.

In recent years, there has been an increased focus on the importance of diversity, equity, and inclusion (DEI), and funding agencies are placing an ever-increasing emphasis on incorporating DEI principles throughout organizational structures and outcomes (see *Broadening Participation in STEM*, n.d.). In organizations that explicitly identify DEI issues as central to their specific goals and outcomes, one might expect to find DEI principles equally present within their leadership structures. However, the default approaches to leadership continue to reflect historical and hierarchical power structures (Schmitz, 2021; Senge et al., 2015), despite evidence that such approaches are ineffective (Dugan, 2017). Within academia, leadership in DEI-focused organizations and collaborations can be characterized by the tension between the hierarchical structures that continue to persist in academia generally and especially in grant-funded projects, and the values, goals, and structures that align with equity and inclusion work. Examining this tension, understanding its divergences, interpreting its impact on collaborative dynamics, and making sense of how academic leadership in DEI work can successfully evolve is critical to the overall advancement of DEI work. Effectively centering DEI principles necessitates an intentional shift toward inclusive leadership models. However, little guidance exists to support organizations and leaders as they undergo this challenging process.

The purpose of this paper is to advance our understanding of leadership structures in the context of a large-scale, collaborative organization focused on increasing DEI in STEM disciplines within higher education. Specifically, this paper presents a case study of *the Alliance*^1^—a federally funded initiative that formed as a DEI-focused collaboration between six organizations, which came together to join their related research and implementation agendas to form a single collaborative alliance. Following a period of turbulence exacerbated by the incongruence of their DEI principles and initial hierarchical leadership model, the Alliance underwent a process of introspective leadership evaluation and evolution to move the Alliance away from hierarchical leadership toward inclusive leadership in line with their DEI values and research agenda. Our inquiry into the Alliance leadership evolution is guided by two research questions:

1. How have the Alliance leadership structures developed and evolved over the life of the large-scale collaborative alliance?
2. How have DEI principles and collaborative frameworks (i.e., collective impact) affected leadership structures and decision-making within the Alliance?

## Theoretical Framework

Traditionally, leadership has been envisioned and enacted as a top-down power structure in which superiors wield power and control over their subordinates—leaders lead while others follow (Pearce & Conger, 2003). This hierarchical leadership structure reinforces positionality and power dynamics that are at odds with the fundamental values and desired outcomes of equity-focused change work (Dugan, 2017). As Allen et al. (2010) explain, “The patterns of hierarchical leadership that served us in the past are not well suited to the global complexity, rapid change, interdependency, and multifaceted challenges” of the 21st century (p. 248). However, decades of socialized practices perpetuate these traditional forms of hierarchical leadership (Hill et al., 2018) and funding agencies continue to emphasize top-down approaches with a single principal investigator (PI) at the helm (see NSF, 2021).

Alternatively, inclusive approaches support collaborative, empowering, and authentic leadership that centers and promotes DEI principles, within both the organization and the research agenda. Below we explore the three central frameworks that shaped the initiation, development, and evolution of leadership within the Alliance: collective impact (CI), shared leadership, and systems leadership. In their funding proposal, the Alliance identified CI as a central element for operationalizing their leadership structure. Likewise, we employ CI as the framework guiding our analysis of leadership within the Alliance. As we will show, CI as a framework does not specifically guide leadership structures or identify key elements for inclusive leadership. As a result, the Alliance struggled to establish approaches to inclusive leadership. Next, we explore shared leadership and systems leadership as the two most influential frameworks that guided the evolutionary process of leadership in the Alliance. Unlike CI, these frameworks specifically address key elements of enacting inclusive leadership, which provided additional guidance as the Alliance sought to reestablish leadership structures.

### Collective Impact and Leadership

According to Kania and Kramer (2011), CI is “the commitment of a group of important actors from different sectors to a common agenda for solving a specific social problem” (p. 36). CI centers five specific conditions: common agenda, shared measurement systems, mutually reinforcing activities, continuous communication, and backbone support organizations. A *common agenda* ensures that participants have a shared vision for change, a common understanding of the problem, and a joint approach to solving it. *Shared measurement systems* ensure agreement on how success will be measured and reported, and coordinating *mutually reinforcing activities* allows each participant to engage in different activities within their wheelhouse in a way that supports the actions of other participants. Participants build trust through *continuous communication*, which includes facilitating frequent meetings that allow participants to learn and problem-solve together. Finally, a *backbone support organization* is a separate entity that provides infrastructure for collective impact, especially collaborative decision-making through facilitation, communication support, data collection and reporting, and handling logistical and administrative details.

#### Challenges of Collective Impact Leadership

Although CI was designed to be a central framework of the Alliance, as it is in other DEI-focused alliances, the theory itself lacks a clear vision for leadership. In a workshop for new alliances, the National Science Foundation (NSF) noted the need to “cultivate leaders with unique system leadership skills” within CI work (NSF presentation, September 2021). While NSF promotes the broader notion of a “collaborative change framework,” which includes collective impact, they did not, however, identify specific system leadership skills or elaborate on how system leadership can, or should, be incorporated into the CI framework. The CI framework itself requires a break from traditional hierarchical leadership models. As Hanleybrown et al. (2012) explain, CI work “requires a very special type of leader…who is passionately focused on solving a problem but willing to let the participants figure out the answers for themselves, rather than promoting his or her particular point of view” (p. 3). Achieving this form of leadership has been challenging. In their review of 25 CI initiatives, Spark Policy Institute and ORS Impact (2018) found three central leadership challenges: high turnover, a lack of individual effectiveness, and a lack of diversity in leadership. Even within a CI framework, successfully implementing collaborative leadership models remains challenging and current theories lack guidance.

In evaluating their collaborative research partnership, Hill et al. (2018) found that a collective impact approach requires effective, inclusive, and dynamic leadership. In their study, project leaders were more accustomed to top-driven PI structures and hence struggled to establish structures for collective decision-making as well as an environment that welcomed and valued contributions from individuals in all levels of the project. However, noting concerns with hierarchical leadership structures and processes, they found that project leaders needed to “conceptualize and build a leadership structure that empowers project members to achieve collective impact” (Hill et al., 2018, p. 20). Such leadership guidance and empowerment were neither inherent in the CI framework put forward by NSF, nor adapted by the Alliance.

#### Centering Equity in Collective Impact

Recently, Kania et al. (2022) expanded their model of CI to center equity, redefining CI as “a network of community members, organizations, and institutions that advance equity by learning together, aligning, and integrating their actions to achieve population and systems-level change” (p. 38). They outline five strategies for centering equity in CI: 1) ground the work in data and context, and target solutions; 2) focus on systems change in addition to programs and services; 3) shift power within the collaborative; 4) listen to and act with community; 5) build equitable leadership and accountability (p. 41). These strategies recontextualize CI within an equitable vision for collaborative systems change—a vision which became a driving factor for leadership evolution within the Alliance. As Kania et al. (2022) explain, “Too often, we focus on diversity to change who sits at the table without changing the underlying dynamics of decisions made at the table by shifting culture and power. Equitable results require more equitable decision-making tables” (p. 43). CI requires commitment to and actions that embrace equitable processes within collaborative initiatives. In addition to shifting power within the collaborative, Kania et al. (2022) promote a distributed form of leadership, which extends to every level of the collaboration. To this end, projects must intentionally create a backbone team that reflects the diversity of the group being served (in our case, people marginalized in STEM) and hold people in positions of power accountable to growing in their own equity work. This shift requires that leaders “do personal, deep introspection to understand their own contributions to the status quo… Structurally, maintaining accountability for equity leadership can be difficult because collective impact is a nonhierarchical approach” (Kania et al., 2022, p. 45). While this new perspective on CI takes the important step of intentionally centering equity within the process of CI work, it stops short of providing guidance on how CI collaborations can achieve equitable forms of shared leadership and disrupt hierarchical leadership structures.

### Shared Leadership

While traditional models of leadership focus on downward influence, shared leadership is defined as “a dynamic, interactive influence process among individuals in groups for which the objective is to lead one another to the achievement of group or organizational goals or both” (Pearce and Conger, 2003, p. 167). Shared leadership impacts organizational culture by emphasizing collective and collaborative practices, as opposed to traditional executive director culture in which all major decisions are made or approved by an executive director (Rothieaux, 2015). Shared leadership can be characterized by four common themes: distributed leadership, decentralized decision-making, recognition and incorporation of diverse perspectives, and an understanding that collective input, deliberation, and decision-making improves the quality and effectiveness of decision-making and implementation (Rothieaux, 2015). Enacting shared leadership relies on establishing a sense of empowerment, trust, and common purpose among all levels of the organization. Ultimately, shared leadership “rests on a commitment by top leaders to share power and foster a climate of trust, safety, fairness, and support” (Freund, 2017, p. 18).

Shared leadership can provide a framework for developing inclusive leadership that aligns with DEI principles. However, organizations seeking to implement shared leadership will likely face what Fletcher and Käufer (2003) identify as three paradoxes. First, it is paradoxical that hierarchical leaders are charged with creating less hierarchical organizations. Hierarchical leadership structures continue to prevail in academic research initiatives, such as those funded by federal agencies (e.g., NSF) that require the use of a single PI in their grant applications (NSF, 2021). PI models reflect and preserve hierarchical approaches to leadership.

Second, it is paradoxical that the rhetoric of shared leadership focuses on collaboration and collective learning, while the imagery of what it means to be a leader perpetuates images of heroic individualism. Fletcher and Käufer (2003) found that shared leadership practices are “often linked to interpersonal attributes and noted as personality characteristics rather than leadership skills,” which is especially true for women (p. 26). Shared leadership is a “participatory process” in which leaders must remain committed to empowering and supporting their team, learning new ways of leading, and letting go of control (Freund, 2017, p. 18). Without commitment at the highest levels, shared leadership fails to take root.

Third, it is paradoxical that leadership roles continue to center the myth of meritocracy and individual achievement, even as they espouse the language of collaborative action. While the language of empowerment is attractive and democratic, failure to deliver on promises of shared leadership “can lead to even greater cynicism about leadership, alienation, and abdication of moral responsibility” (Ciulla, 2010, p. 196). As Ciulla (2010) explains, “when you really empower people, you don’t just empower them to agree with you” (p. 207). Leaders must rely on responsibility, trust, respect, and loyalty to move beyond rhetoric and generate authentic empowerment. To overcome these paradoxes or tensions, Fletcher and Käufer (2003) propose rethinking shared leadership from a relational perspective, specifically situating these processes within the societal contexts of gender and power. Addressing these paradoxes of shared leadership are central to the DEI-focused work of organizations seeking to apply DEI principles to their internal processes as well as the product of their work.

### System Leadership

Theories of system leadership also focus on establishing a collective and collaborative approach to leadership. In their examination of system leadership, Senge et al. (2015) found that system leaders exhibit three core capabilities, which include the ability to see the larger system, foster reflection and generative conversations, and shift the collective focus from reactive problem-solving to co-creating the future. In system leadership, leaders take on the role of facilitator rather than sole decision maker. Senge et al. (2015) also identify three gateways to becoming a system leader. First, a system leader must redirect attention inward, recognizing that the “problem” is internal as well as external: “Real change starts with recognizing that we are part of the systems we seek to change” (p. 29). Such recognition is critical to DEI-focused work, especially alliances with leaders from historically privileged groups (i.e., white and male). Second, a system leader must create opportunities for change: “System leaders work to create the space where people living with the problem can come together to tell the truth, think more deeply about what is really happening, explore options beyond popular thinking, and search for higher leverage changes through progressive cycles of action and reflection and learning over time” (Senge et al., 2015, p. 30). System leadership requires creating accessible decision-making arenas and empowering participants at every level to engage in the decision-making process. Third, they emphasize the importance of practice in becoming a system leader, specifically through implementing tools with regularity and discipline. These tools include system mapping, tools for fostering reflection and generative conversations, and building the capacity to shift from reacting to co-creating. The Alliance employed these tools during their evolutionary process.

## Methodology

This paper examines the Alliance, a large-scale collaborative alliance formed as part of the NSF INCLUDES initiative, a national initiative which sought to broaden participation in STEM fields by increasing participation of female and racially minoritized students—groups which have been historically underrepresented and underserved in STEM education. The Alliance is a collaborative, higher education initiative that seeks to advance DEI reform, which consists of more than 50 partner institutions and organizations from across the United States. While leadership exists at several levels of the Alliance, this paper focuses on leadership at the highest levels, including the principal and co-principal investigators, co-directors, backbone team, and sub-team leadership members.

Data consisted of project documents and semi-structured interviews. Project documents included the grant proposal, meeting agendas, detailed meeting notes, strategic planning documents, co-constructed working documents, visual presentations, Alliance-wide communications, and internal reports. These documents are what Marotzki et al. (2014) refer to as “dynamic data” in that they were developed in interactive contexts. For example, meeting notes often included sections of text copied from the Zoom chat and comments inserted by participants during or asynchronously following the meeting. Documents were downloaded from the shared drive for qualitative analysis, thus becoming “static data” (Marotzki et al., 2014). As part of a larger, IRB-approved research study, semi-structured interviews (*n*=16) were conducted with formal leaders of the Alliance and key representatives from sub-projects within the Alliance and external collaborators to examine collaborative dynamics within the Alliance. Interviews were conducted via Zoom and recorded for transcription.

Data were analyzed through a critical lens informed by applied critical leadership theory (Santamaría & Santamaría, 2012, 2015), critical race theory (Ladson-Billings & Tate, 1995), intersectionality (Crenshaw, 2017), and grounded humanism (Ospina et al., 2012). Analyzing DEI principles within a DEI-focused project necessitates a critical lens to inform our analysis of power structures and equitable leadership. Applying a critical lens allowed us to question the effects of hierarchical leadership against historical inequities and to push back on assumptions of what leadership should look like within DEI-focused research. Dedoose (2021), a web-based, qualitative data analysis tool, was used to code the document and interview datasets. Document data were analyzed using an inductive strategy inspired by grounded theory (Corbin & Strauss, 2008; Glaser & Strauss, 1967). Codes were developed, refined, and applied during two coding rounds by the first author. Interview data were coded using a structural coding approach based on the interview protocol (MacQueen et al., 1998), followed by two rounds of thematic analysis guided by our research questions (Flick, 2013). Lastly, key themes across the second round of coding were identified.

## Findings

The findings from this case study are organized chronologically into five phases, which correspond with the development and evolution of leadership structures within the Alliance: Phase 1) proposed leadership model, phase 2) initial leadership implementation, phase 3) approaching restructuring, phase 4) leadership reorganization, and phase 5) post-reorganization leadership assessment. Figure 1 presents a timeline of leadership structures aligned with the five phases and highlights key Alliance activities in leadership evolution.

**Figure 1.**
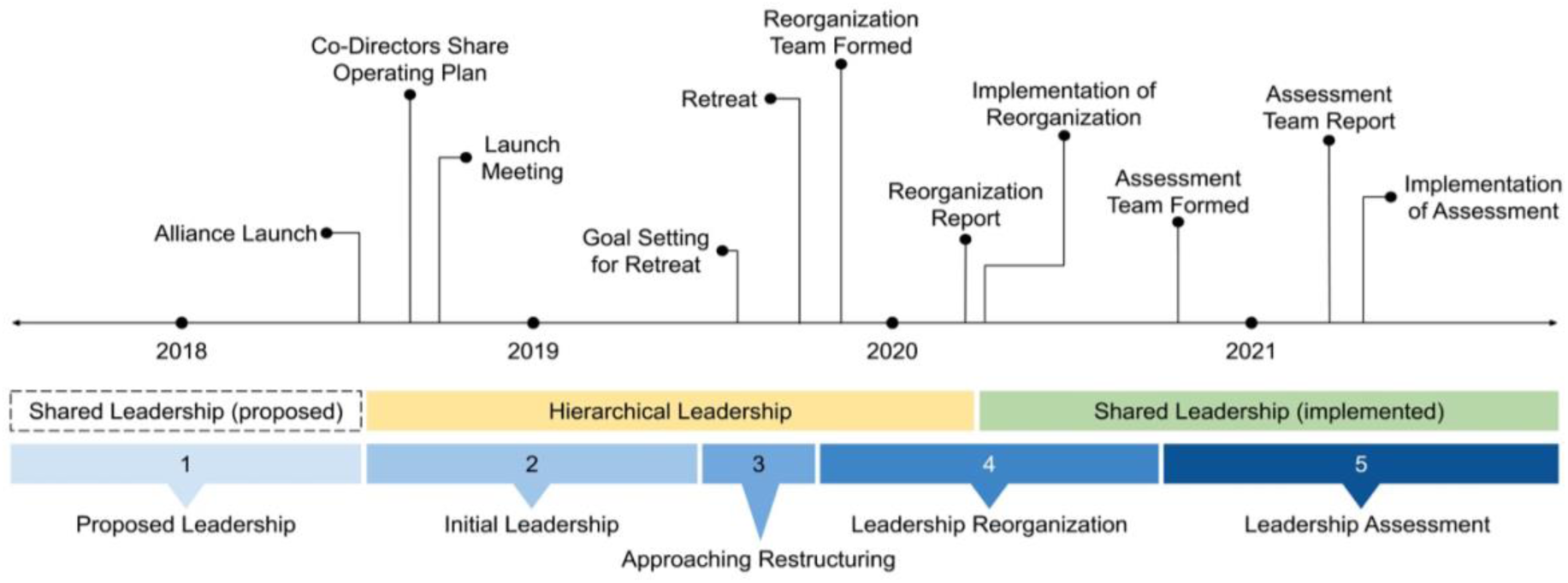
Timeline of Alliance Activities and Leadership Structures

### Phase 1: Proposed Leadership Model

The Alliance formed in response to the NSF INCLUDES Alliance Solicitation (NSF 18-529), which sought to build on the activities of smaller and more localized, ‘launch’ pilot projects by funding the formation of large, collaborative networks. These networks were tasked with developing a vision and strategy for broadening participation in STEM nationwide, contributing to the knowledge base, developing multi-stakeholder partnerships and building infrastructure to support them, establishing a backbone organization to provide communication and networking, and advancing a logic model. Alliances were called on to implement collaborative infrastructure (i.e., shared vision; partnerships; goals and metrics; leadership and communication; sustainability, expansion, and scaling) and collaborative change strategies (e.g., collective impact, network improvement community, participatory action research). According to NSF, approximately 78 percent of the NSF INCLUDES launch pilots reported following the Collective Impact Framework, including the Alliance (NSF INCLUDES Alliance Solicitation 18-529 Information Webinar).

While the CI Framework provides a model for collaborative change (see Kania & Kramer, 2011), it does not provide a leadership framework that is compatible with this approach to collaborative change. However, according to the NSF INCLUDES Alliance Solicitation:

> More than any other element, leadership and communication is particularly important and is what distinguishes NSF INCLUDES from any other program for broadening participation. It is the potential for shared leadership and the building of capacity for leadership and communication across the entire Alliance that provides the glue that will give collaborative infrastructure its ability to function as more than just a collection of organizations. (NSF 18-529, p. 37)

NSF emphasized the importance of leadership, even highlighting the “potential for shared leadership” and communication strategies aligned with collaborative infrastructure. However, they did not offer specific guidance for putting such ideals into practice, leaving each NSF INCLUDES Alliance to develop their own leadership structures.

As the Alliance took shape, so did their proposed leadership structure. They proposed a collaborative leadership model to align with their use of the CI Framework: “The governance system for the Alliance must align and reinforce the CI elements of mutual decision-making, integration of diverse perspectives, and the key role of data, while maintaining clear lines of responsibility necessary to ensure high quality of work, clear direction, accountability, and leadership” (Alliance Proposal). The proposal outlined a collaborative, collective model of leadership based on dynamic governance—an approach to decision-making and governance that allows an organization to manage itself as an organic whole (Buck & Endenburg, 2012). Dynamic governance requires careful implementation planning. When implemented successfully, dynamic governance promotes creative problem solving, accelerates adaptation to change, and engages the energy of all stakeholders. However, dynamic governance can arouse feelings of discomfort for “those not accustomed to sharing the responsibility of difficult decisions” (Buck & Endenburg, 2012, p. 21). According to the Alliance proposal, “Decision-making on overarching issues for the [Alliance] rests with the members as a whole, while significant decisions within sub-teams are made by the team lead(s) and members.” The proposed leadership structure emphasized continuous communication as an important part of providing a “platform for trust to be developed, all voices heard and concerns to be addressed, and ideas to be discussed among groups with different roles” (Alliance Proposal).

### Phase 2: Initial Leadership Implementation

When the Alliance first launched, the proposed leadership structure was relatively flat, with the Leadership Team (LT) working alongside the sub-groups and backbone team. The LT consisted of two co-directors, fourteen sub-group co-leads (two from each of seven sub-groups), and a program manager. In our examination of the data, important questions emerged regarding leadership roles and responsibilities within the Alliance. These questions fell into four key areas, which are often identified as areas of concern in literature on collaborative leadership (see Kania et al., 2021; Schmitz, 2021; Spark Policy Institute & ORS Impact, 2018): shared decision-making, mutual agendas, continual communication, and conflict resolution. Table 1 presents examples of questions regarding shared leadership in these four areas, which were raised multiple times by Alliance members during LT, alliance-wide, and sub-group meetings during the first five months of the Alliance.

**Table 1.** Emerging Areas of Concern

| Area of Concern | Questions Concerning Leadership Roles and Responsibilities |
| --- | --- |
| Shared Decision-Making | • How will shared decisions be made and who will facilitate the shared decision-making process? Who will make the “final” decision?<br>• What role will the co-directors play in shared decision-making? How transparently should decisions be communicated? What happens if the co-directors disagree?<br>• What is the balance between getting things done and slowing down to ensure co-construction and increased engagement? |
| Mutual Agendas | • How will multiple voices be empowered to co-create an agenda?<br>• What will happen when important issues are raised during a meeting that are not in the present agenda?<br>• Who is empowered to change agendas? |
| Continual Communication | • How will the Alliance ensure that “all voices” are heard?<br>• How will diverse voices be heard (i.e., two-way communication) and genuinely incorporated into the Alliance? |
| Conflict Resolution | • What is the process for resolving conflicts within the Alliance, especially conflicts involving a power imbalance?<br>• How should issues and concerns be raised, acknowledged, and responded to within the Alliance?<br>• Who is responsible for addressing conflict? Who is empowered to act?<br>• How does conflict resolution fit within a shared leadership framework? |

*Note.* Questions are a compilation of those raised by Alliance members. These questions were combined and edited for clarity.

Despite the Alliance’s proposed plan for a leadership model that encouraged shared decision-making among Alliance members, the leadership structures and practices implemented during the initial leadership development restructured the leadership to resemble a traditional top-down approach. This shift toward hierarchical decision-making can be seen in the operating plan put forward by the co-directors during the first alliance-wide meeting, in which they detail their leadership approach. LT meetings were the locus of decision-making within the Alliance. While the co-directors note that the “intention has been that Leadership Team meetings should be open to anyone in the Alliance who wishes to attend,” they go on to suggest that LT meetings should remain closed to non-LT members while the LT develops a “mutual understanding that would promote being able to make consensus decisions with each other” (operating plan). They also note that LT members “may need to have a preference” over “others who attend” on “occasions when there isn’t enough time for everyone to speak” (operating plan). Additionally, the co-directors assert that “in those rare times in which agreement is not easily forthcoming, closure still needs to be reached. In such times the co-directors of the project will have to make a final decision” (operating plan). In their critique of the operating plan, one Alliance sub-group co-lead emphasized that “everyone has a voice and is part of collective decision-making” within the Alliance (meeting minutes). They note that the operating plan highlights a “tension between flat structure and hierarchy” and suggests that the leadership structures, particularly those related to communication, were “becoming a broken system” (meeting minutes). Initial Alliance decision-making was situated within the LT, which relied on the co-directors to make final decisions. As one Alliance member commented, “This may reflect the bigger question of who gets to set community norms and rules for decision-making. Is that a function of the LT, or does the LT develop a process for co-creating that with the Alliance members?” (meeting minutes). Additional questions emerged during alliance-wide and sub-group discussions concerning how decision-making will be facilitated and by whom, the role of the co-directors, and how the Alliance will balance collective decision-making with “getting things done” (see Table 1).

The Alliance formed with the common goal of broadening participation in STEM through affecting higher education. Findings suggest the Alliance struggled to narrow and set bi-monthly common agendas in pursuit of their larger goals. The operating plan detailed several additional aspects in terms of top-down leadership, including agenda setting, stating that Alliance members could offer “suggestions for future agenda items.” Data suggest that preset agendas took precedence over emergent concerns, including concerns pertaining to DEI issues within the project itself. As one of the co-directors said regarding LT meetings, “the meeting needs to be well run to keep it moving” (meeting minutes). This product and outcome-oriented focus continued as the Alliance leadership (LT) gathered for the ‘project launch meeting’--their first in-person Alliance-wide LT meeting to mark the official launch of the NSF-funded Alliance.

The collective objectives for the launch meeting were focused on developing a positive culture and community, delineating the processes and practices of the LT, and identifying deliverables and outcomes for the first year of the Alliance (meeting agenda). When the issue of framing and defining DEI mindsets came up during the launch meeting, the co-director suggested that “we should revisit” the issue later (meeting minutes). Later in the meeting, when the topic of defining and unpacking DEI as an alliance-wide activity emerged, an LT member commented that it “seems we should go back to the objectives of the meeting” (meeting minutes). Since such topics were not part of the launch meeting agenda, they were frequently brought up without being fully discussed, instead remaining in the launch meeting “parking lot” at the close of the meeting. The data does not suggest that DEI issues were intentionally or maliciously tabled, but that agendas were upheld with some rigidity in order to maintain focus on the research product rather than internal processes. The data does, however, reveal a clear disconnect between the co-created objectives and the meeting agenda.

Leadership structures influenced the flow of communication. During initial leadership development, LT meetings were the primary source of information and its distribution for the Alliance. Individual LT members were tasked with “communicating with [their] teams about Leadership Team discussions and decisions” (operating plan). This meant alliance-wide communication was filtered through 14 individual LT members, distilling information and complicating communication streams. For some teams, this proved to be a successful form of communication: “I always felt like [my sub-group co-leads] made sure that [our sub-group] was represented, had a voice at the table… There was really open communication between what was happening at the leadership team level and back at the [sub-group] level” (interview). However, other participants reported “misinformation” or miscommunication that occurred when information “didn’t disseminate down” (interview). Data suggest that the initial leadership structure consisted of too many individual streams of communication, which unnecessarily distilled information and limited intra-alliance communication.

### Phase 3: Approaching Restructuring

Issues pertaining to leadership, agendas, decision-making, and DEI continued to grow and, in some cases, fester. The need for conflict resolution first arose during the initial leadership development (see Table 1), but in Spring/Summer 2019 the Alliance determined it was necessary to bring in an external facilitator to help “lower tensions” and “develop guidelines for resolving conflict” (meeting minutes). In preparation for their work with the consultant, the Alliance collaboratively generated a list of goals “to provide a starting point … [and] work on co-constructing possible outcomes” (meeting minutes). The Alliance used an anonymous Google form and a collaborative Google doc to develop their list of goals and outcomes for working with the consultant, to which members asynchronously added their comments, questions, and concerns. Analysis of this document indicates that Alliance members were concerned with the lack of attention given to DEI issues within the alliance leadership structure up to this point. For example, one member noted the need to “develop and apply strategies to more fully elevate all voices and perspectives” within the Alliance, highlighting a common connection of Alliance members’ internal-facing priorities with the Alliance’s external goals to create access for and success in higher education for underrepresented minority students. Another member commented on the need to “unpack power, positionality, and social identity” among members and how they “privilege and/or marginalize” voices within the project. Up to this point, the Alliance had not successfully implemented strategies to ensure the inclusion of all voices and perspectives.

Although facilitating such DEI-focused self-reflection was a primary goal for one of the sub-groups, the Alliance had yet to address these issues internally: “To me, what’s kind of ironic is that where we really struggled was when we start[ed] talking about equity and we turned the mirror on ourselves” (interview). Facilitating critical self-reflection and conflict resolution requires both time and intentionality, which the Alliance had not explicitly planned for in their project design, even as a DEI-focused research collaborative: “I think we perhaps underestimate the extent to which it was difficult and complicated work” (interview). In the collaborative document, members also indicated the need to “support inclusive and brave spaces” that are “free of fear of retaliation, bullying, and harassment”—particularly related to gender, racial, and power dynamics. Several comments in the collaborative Google doc indicate that the Alliance had failed to effectively articulate and communicate roles, responsibilities, expectations (implicit and explicit), decision-making processes, and group norms—central features of collective impact.

Out of respect for the individuals involved in this extremely difficult phase of the Alliance, details of the retreat and work with the external facilitator were excluded from this paper. The retreat represented an intense nadir of conflict seeking resolution for the 20 members of the Alliance who participated (LT members and a small group of core members) and marked a distinct turning point for everyone involved. In the months following the retreat, several members left the Alliance. Ultimately, the Alliance’s work with the consultant was ineffective at addressing these issues. However, examining their process of seeking input highlights the challenges the Alliance was experiencing. Working with the consultant was successful in that it helped them stop “dancing around issues all the time” and instead acknowledge “something’s happening that we need to address” (interview).

### Phase 4: Leadership Reorganization

Following the Fall 2019 retreat, the Alliance established the Reorganization Team (RT), five senior members tasked with “identifying an alternative model for the Alliance’s leadership.” The RT began their work by considering the Alliance’s vision and strategic goals “as a way to prioritize a leadership restructuring that would focus on an optimal organization to achieve these stated goals and aims within a collaborative change systems approach” (meeting minutes). Next, they reimagined leadership possibilities that would support these goals and priorities, which guided their review of the literature and helped them identify salient themes. Following this process, the RT proposed a “shared system leadership” approach for the Alliance, drawing from theories of system leadership (Senge et al., 2015), shared leadership (Pearce & Conger, 2003; Routhieaux, 2015), and adaptive leadership (Heifetz et al., 2004) as described in frameworks above.

Although they initially planned to finish the work in two months, revising the leadership structure of a large-scale collaborative alliance took longer than expected: “It’s hard and complicated work and did not move as quickly as we had hoped initially; we did good work, it just took a longer time to get through” (meeting notes). In January 2020, the RT presented their proposal for leadership reorganization and “facilitated a consensus oriented decision-making process leading to adoption of a shared leadership structure” (annual report). The discussion began asynchronously, with the RT providing a draft of their Revised Leadership Proposal for comments, questions, and conversation. In the proposal, they note that “accountability and communication are essential to achieving a shared systems leadership approach” (reorganization proposal). They went on to ask that all LT members agree to engage in the work and adopt “an approach that reflects a collective, equity-based organizational process” (leadership reorganization proposal). At the next LT meeting, members worked together with the RT to unpack the “tension and anxiety that change produces” (meeting minutes). One RT member reflected on the ways in which “operating within the NSF system” and “having hierarchical experiences” could contribute to anxiety as the Alliance sought to move away from these familiar systems of hierarchical leadership toward a less familiar system of shared leadership. During the discussion, important questions emerged pertaining to the need for a specific decision-making process and communication plan—two of the issues that were central during the initial leadership development and highlighted as challenges from the launch meeting forward. LT members also noted that there were “unresolved issues of trust” that needed to be addressed before trusting the Network (i.e., the proposed LT replacement) to be a decision-making group.

After four months of reflection, research, and revisions, the RT presented a new leadership model, which sought to redistribute project leadership “*throughout* the entire alliance” (document). This new model relied on three Strategic Teams: 1) evaluation team, 2) research team, and 3) an expanded backbone team, which included the new Network team. The Network consisted of the remaining Alliance director and one member from each sub-group, which would most likely be one of the co-leads. Reorganizing the Alliance leadership structures required vulnerability in the face of difficulty and discomfort. As one participant explained:

> What I think has worked is people have been vulnerable and vocal and they’ve, for the most part, stayed with the work even when it got difficult. Even when we had to have difficult conversations and we weren’t on the same page, we’re able to sort of bring that out and kind of process through it. I think that’s helped us build back better or reframe things and make intentional commitments to create a space where we can be, where we can bring out attention, where we can support each other, where somebody actually calls out *okay, hey, wait, I just want to call attention to this* with some intention of trying to get better at working together and coming up with processes. (interview)

This vulnerability and process-level engagement highlights a shift in the Alliance away from the product-oriented drive of top-down decision-making and toward inclusive leadership.

### Phase 5: Post-Reorganization Leadership Assessment

In October 2020, six months after implementation, the Alliance assembled a new team to “conduct a transparent assessment of [their] leadership and organizational evolution that provides opportunities for all Alliance members to provide feedback in a timely and efficient process that results in a more effective alliance” (meeting notes). The Assessment Team (AT) designed and executed a review of the new leadership model through focus groups, anonymous surveys, individual interviews, and review of collective impact survey results (December 2020). Focus group, survey, and interview data were analyzed for emergent themes related to three guiding questions: 1) What changes should be considered to the current organizational structure of the Alliance? 2) What are the current challenges we are facing in leadership and effectiveness of the Alliance, and how might we address them? 3) How well are we engaging members of the Alliance who do not serve in co-lead roles in decision-making, consensus building, and knowledge sharing?

Based on their analysis, the AT put forward a collaborative document to share their initial findings and request input on their suggested solutions (March 2021). This document provided an overview of the background and goals of the AT, their methodological approach, and the results and recommendations based on their strategic analysis of the Alliance leadership structures. The AT invited everyone in the Alliance to review and engage with this asynchronous working document. They also facilitated small group discussions to engage with the working document during an alliance-wide meeting. Finally, having gathered and responded to feedback from across the Alliance, the AT assembled and presented their final report during an alliance-wide meeting (April 2021). Their assessment specifically focussed on the current leadership challenges, efficacy of current structures, and broader engagement of the Alliance.

Regarding the current leadership challenges, the AT put forward three key findings. First, they found that most Alliance members were satisfied with leadership, both broadly and within their teams. Second, communication was perceived as being better, but remained an area of concern for sub-groups. As one respondent put it, “although the Alliance feels more inclusive than it did in the beginning, I hope leaders continue to work on communication (promptness and diversity of voices)” (collaborative document). Third, they found that some Alliance members expressed concern with the number of leadership positions housed within a single organization, which put “a lot of decision-making power and initiative” within that organization (collaborative document).

In examining the efficacy of current structures, the AT put forward four key findings. First, they found that there was substantial confusion among Alliance members regarding the difference between the Network and the Strategic Teams. Second, much like with the former LT, participants reported that the Network “feels like a closed group,” while the Strategic Teams have an “openness [that] makes people feel welcome” (collaborative document). Third, they found that Alliance members were limited by the time and capacity they had to serve on these new teams, leading to a concern that those with more paid time “end up having more voice” (collaborative document). Fourth, they found a lack of clarification regarding the role of backbone within the Alliance.

In examining broader engagement within the Alliance, they found that half of the Alliance members felt unengaged or only partially engaged in decision-making. However, most members reported feeling generally welcomed and listened to within the Alliance.

Based on their findings, the AT put forward three recommendations for further adapting the leadership model. First, they recommended that the Network and the Strategic Team be merged into a single entity with the goal of increasing open and inclusive decision-making. Second, they suggested that co-leads take on a greater role in communicating with their teams to increase the flow of information to team members. Third, they indicated that Alliance should clearly signal decision-making to provide opportunities for input and flatten decision-making structures. The AT put forward recommendations specifically focused on increasing diversity, equity, and inclusion within the Alliance leadership. As one Alliance member explained, “If we’re doing DEI work, you know, there are voices and perspectives in our co-leads that should have kind of equal weight as a co-director” (interview).

## Discussion

The Alliance formed with the best of intentions for applying DEI-focused leadership frameworks. However, the Alliance also formed within the existing hierarchical academic structures and transitioning from proposal to implementation proved challenging, even with many leaders in the Alliance considering themselves to be well-versed in CI practices. The ideals of inclusivity and the realities of existing hierarchies and socialized practices clashed, hindering the Alliance from operationalizing their intended collaborative leadership structures, and likely limiting the realization of their STEM equity goals. For the Alliance, this clash marked the collision of the historical and social structures of academia, the hierarchical organizations from which Alliance members came and from which their practices originated, and their individual identities and previous experiences all with the goals and processes of doing DEI work. Hierarchical leadership structures are deeply intertwined within traditional PI models, and naturally beget hierarchical leaders (Senge et al., 2015). They are a product of culture, rather than the result of strategy (Schmitz, 2021). Such leadership reinforces positionality and power dynamics that are antithetical to the fundamental values and desired outcomes of equity-focused change work (Dugan, 2017). DEI-focused collaborations require a shift in culture and leaders who recognize their own socialized behaviors and are willing to reinvent them.

A chasm exists between the intentions and implementations of DEI principles, which collaboratives must successfully navigate if they are to embody the principles they espouse. Although the intentionality was in the Alliance proposal, the reality of engaging in large-scale, DEI-focused collaborative research requires actively engaging in co-constructing spaces, unpacking identity and power, and building specific strategies for collaboration and conflict resolution (Fletcher & Käufer, 2003; Freund, 2017; Senge et al., 2015). During the first two phases of the Alliance, members became so focused on the product of the Alliance (e.g., research findings) that they were unable or unwilling to see the importance of spending time focusing on the process of the work, such as setting aside the hours of time necessary for implementing their proposed shared leadership structures (see Hill et al., 2018). Rather, the Alliance activities and products became nearly the sole focus of Alliance work and non-agenda items were frequently tabled to preserve the flow of meetings and their ‘productivity.’ Product-focused agendas trumped process-focused concerns. Successfully implementing shared leadership approaches requires time, resources, and funding on the front end.

### Collective Impact and the Process of Process

In their examination of Collective Impact (CI), Hill et al. (2018) examine the action and outcome of a research collaborative (i.e., the product), and the process through which this work was being done. In the context of CI, *process* narrowly focuses on the process of creating the product—the outcome of the project. However, the findings of this work indicate that the CI framework should be expanded to encompass the examination of the process of creating, guiding, and adapting the process itself—the *process of process*. For example, *common agenda* as a specific condition of CI necessitates a common understanding of the problem (Kania & Kramer, 2011). For the Alliance, this is a common understanding of the barriers that need to be addressed to improve DEI in STEM education, which speaks to the external product of the Alliance. To apply CI to the *process of process*, the Alliance also needed to develop a *common process agenda* to ensure that participants had a shared vision for change *within* the Alliance, a common understanding of the incompatibility of DEI principles and hierarchical leadership structures, and a joint approach to solving this problem—the internal process of the Alliance.

The other four conditions of CI are similarly reflected in the process through which Alliance leadership evolved. During the leadership reorganization and assessment phases, the Alliance came to an agreement on how success would be measured and reported—a *shared measurements system*. The new leadership structure established *mutually reinforcing activities*, engaging Alliance members in collaboratively adapting and improving the leadership structures to better align with Alliance DEI values. Reorganizing the leadership required that the Alliance build/rebuild trust through *continuous communication* regarding the process of leadership reorganization and assessment, as well as including issues of process as agenda items to allow participants to learn and problem-solve together. The Reorganization and Assessment Teams took on the role of process-focused *backbone support* by providing infrastructure for process-focused collective impact through facilitation, communication support, data collection and reporting, and handling logistical details pertaining to the evolving leadership structures.

### Redressing Inequitable Equity Work

The top-down leadership model that took hold in the early phases of the Alliance was neither aligned with CI frameworks nor consistent with the DEI-principles (Dugan, 2017). The Alliance, like other DEI-focused projects, espoused a set of values inherently aligned with DEI work and members connected the values of the work being done with an expectation for how the work should be performed. These values were clear and unequivocal, and espoused by a group of people who were committed to advancing DEI principles. And yet, these principles were not reflected in the ways in which the work was being done. The process did not align with the product. For the Alliance, this dissonance grew in magnitude until the situation became untenable.

Redressing their inequitable approach to equity work resulted in an intentional shift toward inclusive leadership. During the reorganization and assessment phases, the Alliance recommitted to changing the underlying dynamics within the organization by centering DEI principles within their leadership and decision-making processes (Kania et al., 2022). Leaders committed to engaging in the reflexive process, which included “personal, deep introspection to understand their own contributions to the status quo” and maintaining a process of accountability for upholding DEI principles (Kania et al., 2022, p. 45). Inclusive leadership requires a form of *praxis*—an ongoing process of action and reflection focused on disrupting oppressive, inequitable structures (Freire, 1978/2013; Senge et al., 2015). Inclusive leadership praxis is an ongoing process requiring intentionality and commitment.

### Successfully Rebuilding the Plane Mid-flight

Successfully reforming leadership structures is challenging, but possible. It requires time, commitment, resources, and adequate funding to support equitable leadership and decision-making structures that align with DEI values. This case study of the Alliance presents one model for successfully changing leadership structures to uphold DEI principles in the collaborative research process itself. The Alliance went through a process of healing that specifically focused on improving relational interactions across inequitable power differentials situated within the contexts of race, gender, and power (Fletcher & Käufer, 2003). The process through which the Alliance recognized and addressed the impact of inequitable power structures shifted the collective focus toward co-creating an Alliance that embraced the same DEI principles their research espoused (Senge et al., 2015).

The successful transformation occurred because the Alliance never stopped bringing issues to light, even as they were repeatedly set aside in the name of productivity. Voices that were marginalized and ignored were not quiet and they did not go away. These persistent voices required the Alliance to take significant action and devote significant resources to solving the problem, and they eventually succeeded. The transition took more than two years, requiring many dozens of meetings and hundreds of person-hours. The Reorganization Team met bi-weekly for five months to carefully develop an inclusive leadership structure that would uphold the values of the Alliance and support their DEI-focused work in STEM education. They read research on various inclusive leadership approaches, discussed how to implement these approaches within the Alliance, and made a detailed proposal for an entirely new leadership structure. Everyone in the Alliance was invited to participate in discussing, adapting, and approving this plan for implementation. However, this was not the end of the process. Several months after implementing the new leadership structures, the Alliance assembled the Assessment Team to conduct an evidence-based review of the new leadership model and propose additional changes to better support the Alliance.

## Recommendations

Despite the serious challenges that emerged from entrenched hierarchical leadership structures, the Alliance was able to successfully rebuild their leadership mid-flight. However, implementing inclusive leadership structures at the beginning of the project would have been more efficient and the Alliance would have experienced less conflict and greater success with central goals. Based on the successful evolution of the Alliance leadership structures, we propose several recommendations for improving the development and implementation of inclusive leadership structures within a nationwide DEI initiative.

### Prioritize Detailed Planning that Operationalizes Implementation

Transitioning a collaborative organization from initial idea to operationalized implementation requires detailed planning and resources. The Alliance’s plan to implement inclusive leadership structures lacked sufficient details and failed to operationalize the implementation (see Table 1). In general, a project should create a detailed plan to specifically address questions concerning how structures will support and promote DEI principles within the work. How will you include and prioritize diverse voices in decision-making? How will you address challenges related to shared decision-making, agenda-setting, and conflict resolution? Implementation requires adequate time and space to move proposals into active projects, time which needs to be supported and financed by funding agencies (and proposal referees). The Alliance developed their leadership structures within a framework defined by CI and the NSF model for large-scale collaborative leadership. Neither of these models provide a framework that supports the development and implementation of inclusive leadership; the models indicate that leadership should be inclusive, but they fail to provide a framework for *how* this inclusive leadership can be realized. Therefore, it is up to individual collaborations to dedicate the necessary time to have intentional discussions regarding process, discussions that address issues of how their leadership will function, the values their leadership structures will uphold, and how they will uphold DEI principles in both the processes and product of their work. Process development and implementation needs to be a recognized activity, acknowledging the extensive resources required to do this work.

### Develop a Process for Evaluation and Adaptation

The successful reorganization of leadership within the Alliance relied on a cycle of evaluation and adaptation to transition from hierarchical to inclusive leadership structures. Planning versus implementation are vastly different endeavors and developing a successful leadership structure necessitates evaluation and adaptation. Internal assessment needs to be built into the collaborative design as a periodic piece of the process work. As this case study demonstrates, it is easy to default to hierarchical leadership structures when efficacy demands. We recommend that DEI-focused collaborations incorporate a critical, reflexive process from the beginning to analyze power structures and DEI principles within the project. Such a process requires intentional time and effort, but it is an essential piece for engaging in equitable equity work. Developing a midstream process review, such as the process followed by the AT, allows collaborative projects to develop evidence-based, actionable recommendations for implementing a cycle of improvement within their own organization. This process cannot wait for a yearly evaluation. Instead, it requires persistent, critical reflection on how to improve inclusive leadership and uphold DEI principles.

### Integrate Continual Professional Development

DEI work requires critical self-reflection and a continual process of unlearning and learning. Engaging in equity work necessitates holding a mirror up to ourselves, lest we hypocritically engage in inequitable equity work (Kania et al., 2022). Implementing inclusive leadership that upholds DEI principles is no different. Inclusive leadership requires continual commitment, critical self-reflection, and a cycle of inclusive leadership praxis that focuses on identifying and addressing inequitable forms of leadership (Freund, 2017; Kania et al., 2022; Senge et al., 2015). Hierarchical leadership must be intentionally unlearned. Based on this research, we recommend that individuals in leadership positions participate in professional development around self-reflection, shared leadership, and collective impact. Professional development should move beyond individual inquiry, delving into critical reflection around inclusive leadership skills. This work needs to be done in collaboration with experts and other leaders seeking to break away from traditional, hierarchical structures. Inclusive, shared leadership cannot take place without continually engaging in professional development that centers DEI principles and critically examines identity, power, privilege, and positionality.

## Conclusion

Our nation is committed to DEI-focused research and implementation to address fundamental inequities in higher education and society. Achieving success will require the successful operationalization of inclusive leadership within DEI-focused collaborations. The situation of the Alliance is not unique. In fact, this conflict between academic hierarchy and inclusive leadership is likely the norm rather than a rare event within large-scale collaborations. We encourage scholars, ourselves and others, to look more broadly and comparatively across projects at how the clash of hierarchical leadership and inclusive shared leadership is resolved. The more we present, dissect, and make meaning from this dichotomy, the more projects will be willing to invest the time to develop the necessary structures to support this work from the beginning.

## Footnotes

1 Pseudonyms are used throughout.

